# Single cell ATAC-seq identifies broad changes in neuronal abundance and chromatin accessibility in Down Syndrome

**DOI:** 10.1101/561191

**Authors:** Roman Spektor, Jee Won Yang, Seoyeon Lee, Paul D. Soloway

## Abstract

Down Syndrome (DS) is caused by triplication of chr21 and is associated with cognitive impairment, Alzheimer’s Disease, and other developmental alterations. The Ts65Dn mouse model for DS has triplication of sequences syntenic with human chr21, and traits resembling those seen in humans with DS. We performed single-cell combinatorial indexing assay for transposase accessible chromatin using sequencing (sci-ATAC-seq) on cortices of adult Ts65Dn mice and control littermates. Analyses of 13,766 cells revealed 26 classes of cells. The most abundant class of excitatory neurons was reduced by 17% in Ts65Dn mice, and three of the four most common classes of interneurons were increased by 50%. Ts65Dn mice display changes in accessibility at binding motifs for transcription factors that are determinants of neuronal lineage, and others encoded within triplicated regions. These studies define previously uncharacterized cellular and molecular features of DS, and potential mechanisms underlying the condition.

## Introduction

Down Syndrome (DS), affecting one in 800 births in the United States, causes deficits in spatial, long-term, and short-term memory; and impairments in language development and new skill acquisition (reviewed in ^1^). Alzheimer’s Disease is common in DS, with a much earlier age of onset relative to the general population ^2^. Several gross neuroanatomic abnormalities have been described that accompany DS, including reduced volumes of the hippocampus ^3, 4^, dentate gyrus ^5^, parahippocampal gyrus ^5, 6^, cerebrum ^7, 8^, cerebellum ^7^, and amygdala ^3^. There are larger subcortical gray matter volumes reported in DS ^8^, and general cortical dysgenesis, particularly affecting the frontal lobe ^9^.

Accompanying the gross neuroanatomic abnormalities are cellular changes. GABAergic signaling is altered, which may impair synaptic plasticity, as well as learning and memory by altering the balance of excitatory and inhibitory signaling (reviewed in ^10^). In the hippocampus, there is evidence for less cell proliferation, as fewer cells stain positive for Ki67; and there is also evidence for increased apoptosis, as more cells stain positive caspase 3, and display pyknotic nuclei ^5^. Efforts to document changes in specific cell-types have shown that neuronal populations are generally reduced in abundance and density in the hippocampus, including the dentate gyrus, lateral parahippocampal gyrus, entorhinal cortex, presubiculum ^5^, and in cortical layers II and IV ^9^; in contrast, astrocytes are more abundant in human fetal tissue ^5^. Comprehensive assessments of changes in cellular composition of the DS brain have not been possible to date, owing to limitations of the available assays.

Studies of DS in human have been aided by mouse models, notably the widely used Ts65Dn mouse ^11^. This model harbors a reciprocal translocation, and segmental trisomy, leading to triplication of 132 mouse orthologs of the 225 genes found on human chr21 ^12^. Behavioral, anatomical, histological and molecular analyses of Ts65Dn mice identified traits in common with DS in human (reviewed in ^13^). Notably, relative to control littermates, Ts65Dn perform worse in various learning and memory tasks, and mice have structural abnormalities in several brain regions. Additionally, Ts65Dn mice exhibit hypomyelination of neurons, and have slower neocortical action potential transmission ^14^. They also share many of the gene expression changes found in humans with DS ^15^.

Among the mechanisms proposed to explain how chr. 21 trisomy leads to DS is the gene dosage effect model. This posits that genes within triplicated regions are more highly expressed, owing to their increased copy number, and that the elevated expression levels of one or more of the 225 triplicated genes are responsible for DS traits. An alternative, and not mutually exclusive model to the gene dosage effect model is that the triplicated region binds gene regulatory factors that are present in limiting amounts, altering their levels in cells, and the chromatin and expression states of other critical genes outside of the triplicated region. Consistent with the gene dosage model are results from transcript profiling experiments, which have shown that chr. 21 gene expression is elevated for up to 93 transcripts in various tissues from individuals with DS, and 54 transcripts in three mouse models for DS ^15^. Among the transcripts in the triplicated region that are upregulated in DS and mouse models for DS is *TTC3*, an inhibitor of neuronal differentiation ^15^, which may explain the reductions in neuron abundance and density seen in DS. Triplicated genes also include several transcription- and chromatin-regulatory factors (*BRWD1, HMGN1, PRDM15, DNMT3L, USP16, RUNX1, OLIG2, GABPA, ERG* and *ETS2*), although not all these are upregulated in DS. Chromatin state changes, notably DNA methylation, is altered in DS ^16, 17^. These changes may be due to triplication of the DNA methyltransferase, *DNMT3L*, however, cellular composition differences among the tissues compared could also be the source of DNA methylation changes. These intriguing findings, and the fact that transcription- and chromatin-regulatory factors are encoded within the triplicated region of DS, provide motivation to evaluate chromatin changes associated with DS.

In order to characterize the cellular and molecular changes associated with DS in greater detail, and to gain further insights into the mechanisms influencing DS traits, we subjected cortices of adult Ts65Dn mice, and their control littermates to single-cell combinatorial indexing assay for transposase accessible chromatin using sequencing (sci-ATAC-seq). This strategy eliminates the challenges of identifying or purifying specific cell-types using antibodies or other reagents, and instead enables their identification and quantification based on shared molecular features. It also enables the characterization of chromatin state changes associated with DS for the different cell populations identified, as well as the sequence features at domains undergoing changes in DS. The unbiased approach afforded by sci-ATAC-seq provides novel and unprecedented insights into the cellular and molecular features associated with DS.

## Results

### sci-ATAC-seq library preparation, sequencing, and quality control

To prepare sci-ATAC-seq libraries using adult tissue we implemented several modifications embodied in Omni-ATAC-seq ^18^ to snATAC-seq ^19^. These modifications increase coverage at peaks, remove additional cellular debris, and minimize mitochondrial contamination. Briefly, we disaggregated two mouse cortices each from control (2n) and Ts65Dn (Ts) littermate males using Dounce homogenization, and isolated nuclei using density centrifugation to remove myelin debris prior to tagmentation and sorting. We confirmed successful disaggregation and removal of debris via microscopy and during fluorescence activated nuclei sorting (FANS) (Fig. S1a).

The sci-ATAC-seq workflow relies on combinatorial indexing to acquire data from individual nuclei ^20^. In this strategy, nuclei were equally distributed in a 96-well plate (∼4,800 per well) and subjected to tagmentation of Tn5 transposase carrying 96 distinct oligonucleotide barcode combinations ^21^ in Omni-ATAC-seq buffers. After tagmentation, we pooled nuclei from all wells, and used FANS to sort 25 nuclei per-well into a second set of 96-well PCR plates and subjected the DNA to PCR using an additional 96 barcode combinations per PCR plate. This strategy produced 9,216 barcode combinations per PCR plate, containing libraries from 2,400 single nuclei (Fig. 1a).

**Figure 1:**
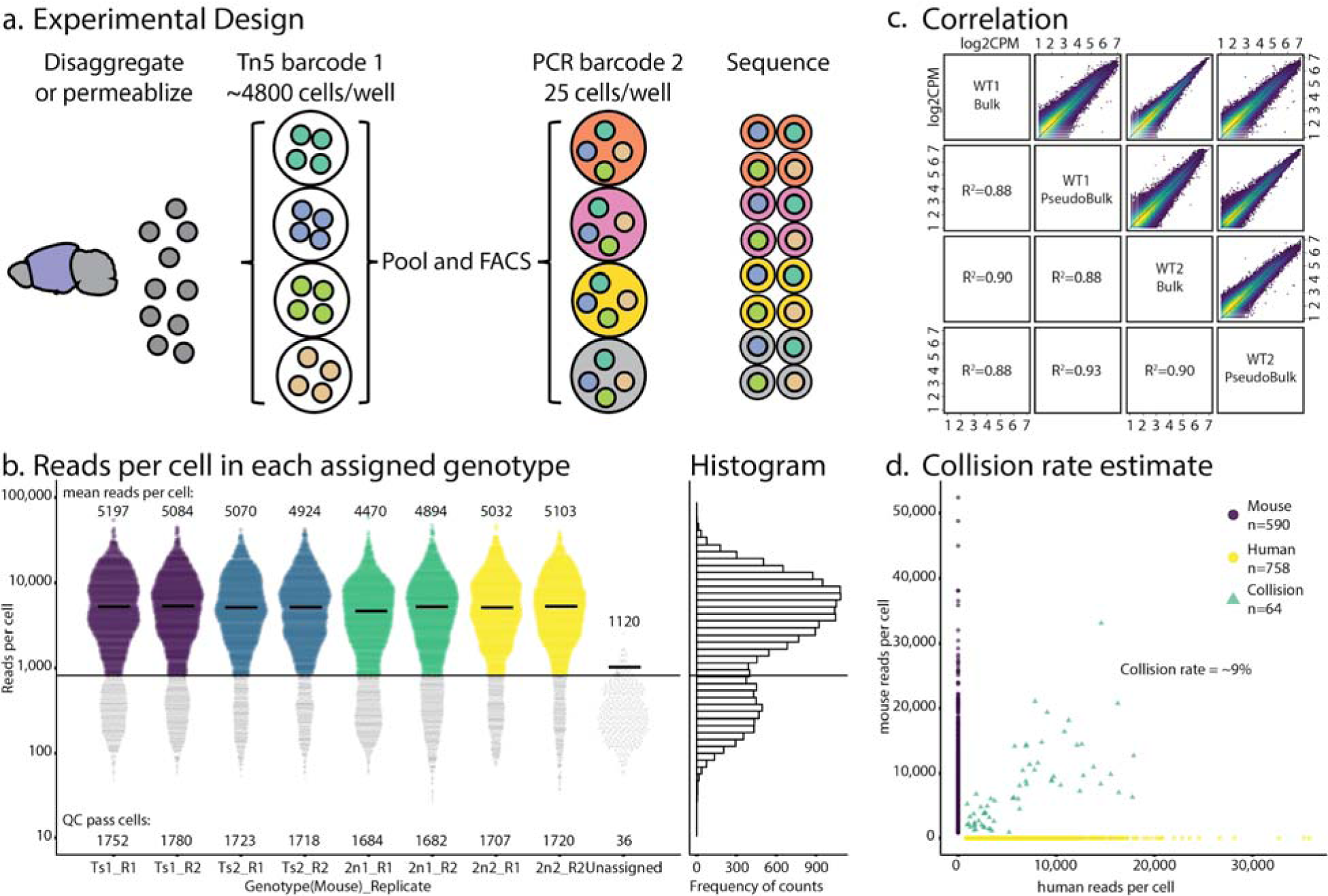
Experimental design and QC. **(a)** The cortex from each brain is disaggregated and nuclei are distributed to 96 wells to be barcoded with Tn5 transposase. Wells are pooled and FACS sorted at 25 nuclei per well. Wells are barcoded via PCR. Sequenced cells are demultiplexed using combinatorial barcodes. **(b)** Reads per QC-passed nuclei per genotype and replicate. Lines denote medians, counts denote means. Line at 820 reads denotes minimum read cutoff. **(c)** log2CPM correlation of each WT brain vs bulk ATAC-seq showing Pearson correlation. **(d)** Collision rate estimates from mixed human and mouse samples.

We pooled DNAs from single cell library plates, and ascertained library fragment size distribution by Agilent Bioanalyzer, observing the size contributions expected from nucleosome DNA length, plus oligonucleotide lengths inserted by Tn5 and PCR (Fig. S1b). Libraries passing this quality measure were quantified by digital PCR (Fig. S1c) ^22^. Sci-seq barcodes incorporated by tagmentation and PCR impart large stretches of identical sequences to each sequenced molecule, which can produce clustering and base calling problems on Illumina sequencers. In order to eliminate this problem, and increase the number of informative reads, we sequenced our libraries using dark cycle chemistry, skipping uninformative and identical bases ^23^. We sequenced a total of 19,200 cells to saturation, with a read duplication rate of ∼80 % (Table S1) on a NextSeq500.

Fig. 1b summarizes our sequencing results, showing the number of unique reads from each cell assayed for each replicate. The aggregate data had a bimodal distribution; we selected 820 reads/cell at the approximate inflection point as our threshold, carrying forward for further analysis the 72 % of cells with reads exceeding our cutoff (13,884 cells). We combined the resulting reads from each assigned sample and mapped them to the mouse genome to identify peaks from the aggregated data, which we define as pseudobulk data. As a further quality control criterion, we evaluated the fraction of reads from single cells that mapped to peaks (FRiP), and used only cells having a FRiP score greater than 20 % (Fig. S1d), as previously used in other sci-ATAC-seq analyses ^24^. 72 %, or 13,766 nuclei pass both quality control criteria and showed expected insert-size distributions (Fig S1e).

To assess the reproducibility of our sci-ATAC-seq libraries, we prepared Omni-ATAC-seq libraries from the same tissues in parallel and made two comparisons. First, we compared data from two replicate bulk libraries, and found them to be highly correlated (R2=0.9 for wild type (WT) cortices, Fig. 1c). Next, we aggregated the single cell data from the two corresponding WT sci-ATAC-seq libraries into pseudobulk datasets, and evaluated how well they correlated with each other, and with the true bulk libraries. R2 values for each comparison were 0.88 or greater (Fig. 1c). We also extended these analyses to the Ts65Dn libraries and found all R2 values were 0.87 or greater (Fig. S2). From these quality control tests, we concluded that our sci-ATAC-seq libraries provided reproducible results, and that the total single cell data accurately reflected the chromatin states observed in the bulk tissues.

As an additional data quality measure, we estimated the frequency with which single cell data were derived from more than one cell. For this test, we made a mixture of mouse and human cells, performed sci-ATAC-seq, and identified the collision rate ^20^, defined as the frequency with which any identified cell barcode had both mouse- and human-specific sequences (Fig. 1d). This value was ∼9 %, within previously reported expectations of ∼12.5 % ^20, 25^.

As a final data quality measure of sequencing coverage, and to confirm proper assignment of samples to barcodes, we created pseudobulk datasets from demultiplexed samples, and performed Copy Number Analysis (Fig. S3a). We expected Ts65Dn pseudobulk data to show evidence for increased copy number of reads within the triplicated region of Mmu16 and Mmu17. This is precisely what we observed, a triplication of Mmu16 spanning chr16:81,000,000-98,200,000 and of a centromeric fragment of Mmu17 at approximately chr17:3,500,001-9,000,000 (Fig. S3b), in agreement with triplicated regions previously reported using Array-CGH ^26^.

### Cell-type identification and validation

We used the sci-ATAC-seq reads from 13,766 cells from both WT and Ts65Dn cortices to perform dimensionality reduction in Monocle to identify a total of 26 cell clusters (Fig. 2a). We assigned the 26 cell clusters identified to the six major cell-types known to be found in cortex: Excitatory neurons (EX), Interneurons (IN), Astrocytes (AC), Oligodendrocytes (OG), Endothelial cells (EC), and Microglia (MG) (Fig. 2b). To do so we aggregated reads from each cluster, preparing cluster- and replicate-specific pseudobulk datasets. We then performed hierarchical clustering of the average relative accessibility at gene bodies of marker genes in each cell-type for each cluster to identify their respective cell-types (Fig. 2c). As confirmation we visually inspected read coverage at marker genes for each cluster (Fig. 2d). We determined that 12 cell clusters represent Excitatory neurons (EX), eight clusters represent Interneurons (IN), three clusters represent Oligodendrocytes (OG), and one cluster each represent Astrocytes (AC), Microglia (MG) and Endothelial cells (EC). To confirm these assignments, we performed the same clustering analysis using previously published bulk ATAC-seq data, generated using FACS-isolated neuronal subtypes ^27, 28^ (Fig. S4).

**Figure 2:**
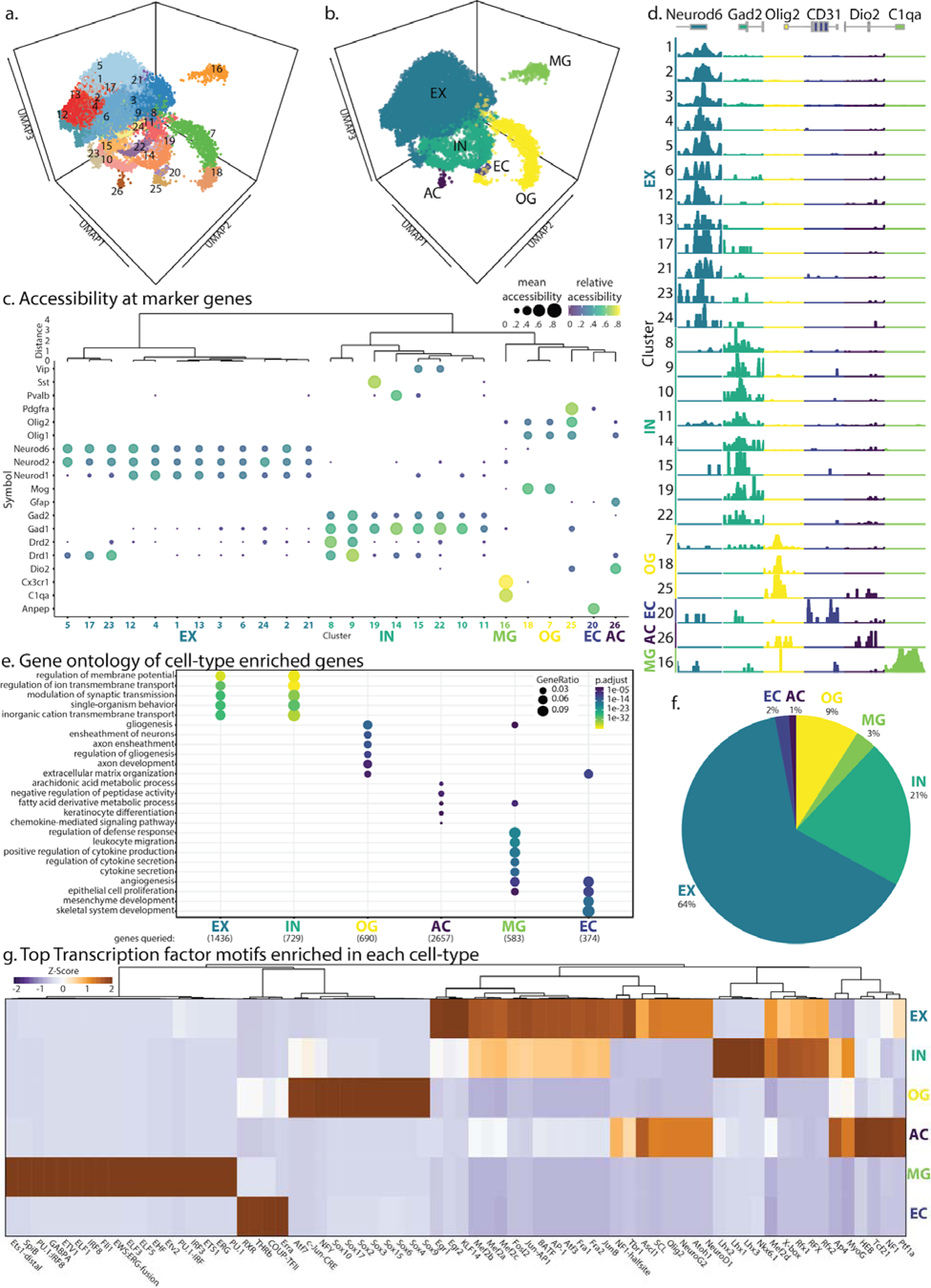
Cell-type identification. **(a)** UMAP of cells showing 26 clusters. **(b)** Clusters from (a) labeled by cell-type **(c)** Clusters are identified using cell-type specific markers. **(d)** Browser shots from clusters in (a) at cell-type specific markers: EX (Neurod6), IN (Gad2), OG (Olig1), EC (CD31), AC (Dio2), and MG (C1qa). **(e)** Genes ontology of genes enriched in each cell-type in (b). **(f)** Cell-type distribution of WT cortex as percent of total. **(g)** Top 20 Transcription factor motifs enriched at peaks in each cell-type (p<1e-20). Cell-type specific transcription factors are color coded.

To classify cells in our EX population, we looked at layer-specific markers. Clusters EXc1, EXc2, EXc4, EXc12, EXc13, and EXc23 are accessible at Cux1, Cux2, and Satb2, localizing to cortical layers II/III/IV ^29^. Cluster EXc3, EXc5, EXc6, EXc17, EXc21, and EXc24 are accessible at Bcl11b while being inaccessible at layers II/III/IV markers indicating their localization to layers V/VI; of these, EXc5 and EXc17 are both more accessible at Foxp2, and Pcp4, indicating their localization to layer V ^30^. In our IN population, we assigned cell-types based on their accessibility at interneuron marker genes. INc8 and INc9 are dopamine D2 and dopamine D1 receptor accessible medium spiny neurons, respectively. Cluster INc14 contains Parvalbumin accessible interneurons, INc19 contain Somatostatin accessible interneurons, and INc15, and INc22 contain Vip accessible interneurons. We were not able to clearly identify the remaining two interneuron clusters; INc10, a class hyper-accessible at Frmd7, Sp8, Vipr2, Etv1, and Ntsr1 which appears to resemble Int16 in Zeisel et al ^31^, possibly localizing to layers V/VI of the cortex ^32^, and INc11 as a class accessible at Lhx8, Zic4, and Myo3a. Of our three oligodendrocyte populations, OGc25 contained oligodendrocyte progenitor cells accessible at Pdgfra while inaccessible at Mog.

To further confirm our cell-type assignments we identified genic regions and their 2kbp flanks that are hyper-accessible in each cell-type, and cluster relative to all other cell-types (Fig. S5a, S5b) and clusters (Fig. S5c-e). Gene Ontology (GO) analysis at hyper-accessible gene bodies in each cell-type showed findings were consistent with expectations for these cell-types (Fig 2e), providing further confidence in our assignment of cells comprising the 26 cell clusters to their corresponding six cell-types. Neurons showed GO enrichment for genes enriched in categories such as regulation of membrane potential, and modulation of synaptic transmission. Oligodendrocytes showed GO enrichment for escheatment of neurons and gliogenesis. Astrocytes showed less indicative GO categories such as arachidonic acid metabolic processes and Keratinocyte differentiation. Microglia showed enrichment in genes involved in cytokine production and leukocyte migration. Endothelial cells showed enrichment for genes involved in angiogenesis and epithelial cell proliferation.

Our assignments match previous reports quantifying cell-types in the cortex ^19^. We observed most cells in WT mice to be excitatory neurons (64 %), followed by interneurons (21 %), oligodendrocytes (9 %), microglia (3 %), endothelial cells (2 %), and astrocytes (1 %) (Fig. 2f). For clarity, we have prefixed all clusters to their assigned cell-types (Table S2a).

We then identified transcription factor (TF) motifs found at peaks of open chromatin that were enriched in each cell-type, relative to others. In EX population, hyper-accessible peaks were enriched for binding sites for TFs contributing to excitatory neuron specification such as NeuroD1 and NeuroG2. Similarly, they were enriched for Tbr1 motifs, a factor important to neuronal migration and axonal projection ^33^. Hyper-accessible peaks in our IN population were enriched Lhx3 motifs, a factor known to have a role in the development of interneurons ^34^ (Fig. 2g). We observed an enrichment of Sox-family TFs in OGs and ETS-family transcription factors in MG populations. In ACs we observed an enrichment of NF1 sites, a regulator of astrocyte proliferation ^35^. Within our ECs we observed an enrichment of COUP-TFII motifs, a regulator of endothelial identity ^36^. Such findings provided additional confidence in our assignments of cell clusters to known cell-types.

### Changes in cell-type representation in Down Syndrome

Having identified 26 cell clusters in adult cortices, and the six cell-types to which they belong, we asked how the Ts65Dn triplication affected these populations. We first evaluated cell abundances. Cortices of DS animals had significantly more interneurons; and specifically, among the eight cell clusters comprising interneurons, five were significantly larger in Ts65Dn animals (Fig. 3a, Fig. 3b). This finding agrees with reports that interneurons are more abundant in DS ^37, 38^. Specifically, we observed a large change in INc8 and INc9, which are Drd2 and Drd1 positive medium spiny neurons, respectively, and INc10 and INc11. We did not observe a statistically significant change in Parvalbumin positive (INc14) or Somatostatin positive (INc19) interneurons (Fig. 3b).

**Figure 3:**
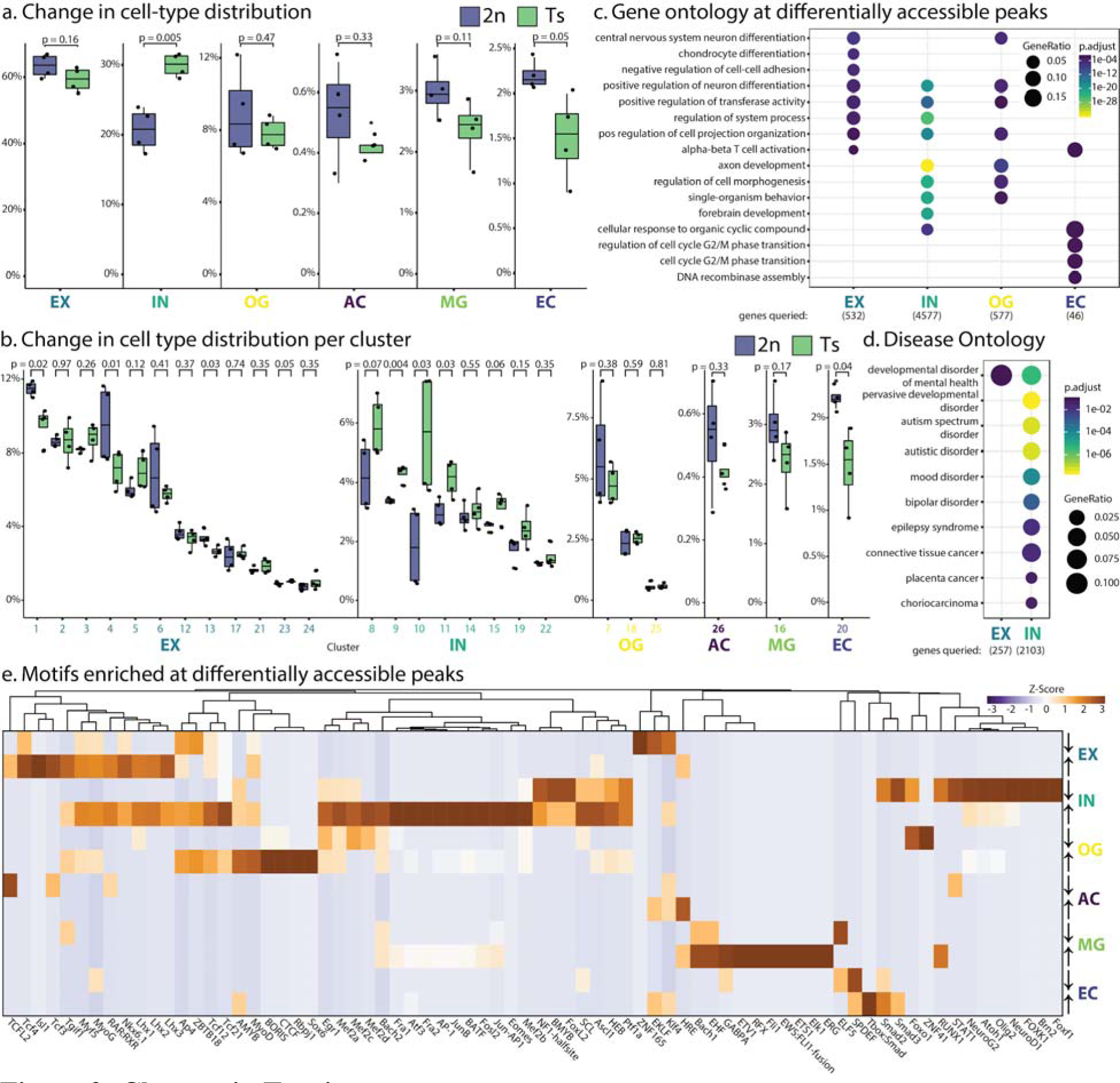
Changes in Ts mice: **(a)** Changes in cell-type (top) and **(b)** cell cluster (bottom) abundances in WT (blue) and Ts (green) mice. P values indicate Student’s t-test. **(c)** Gene Ontology at genes nearest to differentially accessible peaks (p<0.05, FC>1.5) in Ts mice. **(d)** Significantly enriched Disease Ontology in Ts mice at genes from b. **(e)** Top 20 transcription factor motifs in Hyper-accessible (up) or Hypo-accessible (down) peaks in Ts mice. Values denote Z-scores.

Although the change in abundance of excitatory neurons was not statistically significant, among the 12 excitatory neuron clusters, three had significantly fewer cells in the Ts65Dn animals, including the most populous of the 12 clusters, and one had significantly more cells. Each of these clusters appears to localize to layers II/III/IV. This corresponds with known delays in the development of the cortex in prenatal Ts65Dn mice; though this change has not been measured in as great a detail in adults ^39^. Interestingly, the single increased EX cluster showed the highest relative accessibility at Synj1, which regulates levels of membrane phosphatidylinositol-4,5-bisphosphate, and may affect synaptic transmission ^40^. Beside these changes in neuronal populations, we observed significantly fewer endothelial cells than their WT control littermates (Fig. 3a, 3b), a previously undocumented change that may influence impaired endothelial function reported in Down Syndrome patients ^41^.

In addition to performing cell clustering in Monocle, we used a TF-IDF-based pipeline ^25^, identifying 25 cell clusters (Fig. S6). When we assigned these to the six cell-types using the above strategy, we found 92.5 % agreement in our identified cell-types and cell-types identities in clusters called using TF-IDF.

### Gene ontology at differentially-accessible peaks

In order to assess biological processes changed in Ts65Dn mice, we performed Gene Ontology analysis on genes nearest to differentially-accessible peaks within each cell-type (Fig 3c) and cluster, (Fig. S7a) relative to WT mice. In our EX clusters, we observed significant enrichment in ontologies such as CNS development, cell-to-cell adhesion, neuron differentiation, and projection organization. In our IN cell-type, we saw enrichment for neuron differentiation, axon development, and cell morphogenesis, forebrain development, and response to cyclic compounds. In our OG cell-type we observed enrichment in CNS differentiation, adhesion, and cell morphogenesis. In endothelial cells, we saw enrichment of genes involved in the regulation of cell-cycle and phase transition; specifically, genes within these functional categories are hypo-accessible in Ts65Dn, which is likely to correlate with their diminished expression. We observed no enrichment of terms in astrocytes and microglia.

In order to further understand the contribution of differentially accessible peaks within each cell-type to pathology, we performed Disease Ontology analysis on human homologues of genes nearest to differentially-accessible peaks in each cell-type (Figure 3d) and cluster (Fig. S7b). We observed genes nearest to differentially accessible peaks in neuronal clusters were enriched for mood and developmental disorders and epilepsy. Furthermore, in Ts65Dn mice we observe an increased representation of epilepsy-related genes, mood-disorder, and developmental-disorder-related genes in our IN cell-types, with a slight enrichment for genes related to developmental disorders of mental health in our EX cell-type.

### Transcription factor motifs at differentially-accessible peaks

In addition to these measures of cell abundance and ontology analyses, we identified TF binding motifs at differentially accessible peaks between Ts65Dn mice and their WT littermates for each of the six cell-types (Fig 3e) and 26 clusters (Fig. S8). Our first expectation was that we would observe increased accessibility in Ts65Dn mice at motifs associated with triplicated transcription factors (ERG, GABPA Runx1, and Bach1). Indeed, these TFs were among the top 20 enriched TF motifs present at hyper-accessible sites in microglia. Of the 91 hyper-accessible peaks in Ts65Dn microglia, these transcription factors account for 31 %, 24 %, 20 %, and 4.4 %, of peaks observed, respectively.

The most radical change at hypo-accessible peaks interneurons of Ts65Dn mice occurred at bHLH-family TFs (NeuroD1, Olig2, Atoh1, NeuroG2, Tcf21), which have similar motifs, and dictate neuronal differentiation paths. Specifically, we observed enrichment of Olig2 motifs, which is encoded in the triplicated region, at both hyper- and hypo-accessible peaks in Ts65Dn interneurons. Complementary to this, we observed hyper-accessibility at bZIP-family TFs (AP-1, Fra1, JunB, ATF3, Fra2, BATF, Fosl2) in Ts65Dn interneurons; these factors have been shown to be highly active in interneurons after hippocampal seizures ^42^. Outside of this enrichment in interneurons, we observed increased accessibility at motifs for MADS-family TFs (Mef2a, Mef2b, Mef2c), which have been shown to specify neural differentiation and are implicated in neurodevelopmental disorders ^43–45^. Similarly, in Ts65Dn interneurons, we observed an increase in accessibility at motifs matching Eomes, a regulator of neurogenesis ^46^. Overall these results suggest to us a large change in cell-lineage commitment along with potentially aberrant localization of Interneurons.

We observed fewer significant changes outside of Interneurons, the most affected cell-type in Ts65Dn mice. Within excitatory neurons there was an enrichment in Homeobox-family TFs (Lhx1, Lhx2, Lhx3, Isl1, Nkx6.1) at hyper-accessible sites, many of which are involved in interneuron/motor neuron differentiation ^47–49^, and bHLH-family TFs (Myf5, MyoD, MyoG), which have been demonstrated to inhibit neuronal differentiation ^50^.

In oligodendrocytes, we observed an enrichment of CTCF/BORIS motifs at hyper-accessible peaks. CTCF has been implicated as a marker of oligodendrocyte differentiation ^51^. This orthogonally matches GO enrichment at terms relating to differentiation in oligodendrocytes. Defects in oligodendrocytes have previously been seen in Down Syndrome ^14^, both *in vivo* and when cells are differentiated *in vitro*. Concordantly, we see bHLH-family TF motifs (MyoD, Tcf21, Ap3, ZBTB18, NeuroG2, Olig2) at hyper-accessible peaks in Ts65Dn mice. Interestingly, we noted no change in either class of oligodendrocytes (OGc7, OGc18) or oligodendrocyte progenitors (OGc25).

## Discussion

Single cell sequencing has become an increasingly powerful method for measuring disease-associated changes in cell populations, and their chromatin states. In this study, we performed a comprehensive pairwise comparison of chromatin accessibility in the Ts65Dn mouse model for Down Syndrome, and their wild-type littermates using sci-ATAC-seq. From 13,766 randomly sampled cells that met quality control criteria, we identified 26 discernible clusters representing distinct cell-types in the brain cortex, and quantitative changes in cell composition in Ts65Dn mice. Additionally, we identified sites of altered chromatin accessibility caused by the Ts65Dn triplication. Because sci-ATAC-seq requires no cell purification, our identification of cell-types was unbiased, and provides a high resolution of cellular and chromatin state changes in Down Syndrome.

Consistent with previous studies, we observed a broad increase in the number of interneurons in Ts65Dn mice. Specifically, we observed that dopamine-receptor expressing interneurons were affected disproportionately relative to other clusters. Interestingly, our data falls in closer agreement to the studies of Hernandez-Gonzalez et al ^38^, which used similarly aged 4-month-old adult mice, than to those of Chakrabarti et al ^37^, which observed defects in Somatostatin+ interneurons in 1-4-week-old mice. Our data identified no changes in Somatostatin+ and Parvalbumin+ interneurons in adult Ts65Dn mice.

The increase in interneurons was accompanied by a significant decrease in abundance of four of the 12 classes of excitatory neurons we detected, most probably originating from the upper layers of the cortex. These findings agree with other reports that in Ts65Dn mice, there is an imbalance of inter, and excitatory neurons, with the former being elevated, and the latter being diminished ^37^. Such cellular changes are likely to contribute to cognitive deficits associated with Down Syndrome in Ts65Dn mice, as maternal choline supplementation during gestation and lactation-a treatment that partially ameliorate the deficits - also partially normalize cell population changes ^52^.

In addition to these changes in cell abundances, we identified chromatin domains with altered accessibility in Ts65Dn mice relative to their wild-type controls. Transcription factor binding motifs present at those domains were consistent with previous assertions that increased gene dosage of transcription factors within the triplicated region play a role in the mis-regulation in Down syndrome ^37, 53^. Specifically, we observed that sites of altered chromatin accessibility in Ts65Dn microglia were enriched for binding motifs for RUNX1, GABPA, and ERG; and in oligodendrocytes, there was an enrichment of OLIG2 motifs at differentially accessible peaks.

Besides these motifs for transcription factors encoded by the triplicated regions, we observed that CTCF sites were hyper-accessible in oligodendrocyte chromatin from Ts65Dn mice, implying a subtle differentiation defect. Although we did not find changes in OG abundance between Ts65Dn mice and their WT littermates, the altered accessibility at these sites is likely to impart functional changes in cellular behavior. Our lack of statistically significant changes to any of the three oligodendrocyte clusters we observed contrasts with the results reported by Olmos-Serrano et al. However, those studies characterized 1-week old mice, whereas we used 10-week old adult mice in our study. We infer that for these cell populations, delayed differentiation is occurring in Ts65Dn mice, and this may be one of the functional changes caused by altered chromatin accessibility.

Our inferences regarding changes in low abundance cell-types such as endothelial cells, astrocytes, microglia, and minor clusters comprising excitatory and interneurons are limited by the number of cells we analyzed (13,766). More definitive conclusions will require profiling of larger cell numbers. Furthermore, by characterizing mice at a single time point, we cannot define when during development regulatory changes, or changes in cell abundance occur. That will require analysis of mice at different ages. We anticipate that analyses of other brain domains, including hippocampus for example, will reveal additional cellular and molecular changes associated with Down Syndrome. A final challenge is discerning which of the changes detectable by sci-ATAC-seq are important to Down Syndrome traits. This can be facilitated by characterizing brains of Ts65Dn mice whose functional deficits are partially suppressed by perturbations such as maternal choline supplementation ^52, 54–56^, transcription factor dosage compensation ^37^, and drug treatments ^57^.

## Acknowledgements

Jennifer D. Mosher, Ann E. Tate, Jeff C. Mattison, and Peter A. Schweitzer from the Cornell University Biotechnology Resource Center (BRC) for genomic sequencing. Cornell CARE staff. Andrew Adey for sharing their CPT-seq dark cycle chemistry. Danea F. Rebolini at Illumina for modification Adey lab dark cycle chemistry. Andrew Grimson for the gift of A549 and Hepa1-6 cells.

## Data availability

Data have been submitted to NCBI Gene Expression Omnibus; no accession number has been provided yet.

## Methods

### Mice

B6EiC3Sn.BLiA-Ts(1716)65Dn/DnJ (Jackson #001924) mice were purchased from Jackson and crossed to B6EiC3Sn.BLiAF1/J (Jackson #003647) for maintenance. Mice used for sci-ATAC-seq were crossed to B6.Cg-Tg(Thy1-YFP)16Jrs/J (Jackson #003709) to express YFP in excitatory neurons for future analysis (unpublished data). Genotyping PCR was performed on 50 ng genomic DNA using oligos from Table S3 from for 40 cycles using GoTaq (Promega #M3001) (94°C 2min, 40 cycles of: [94°C 30sec, 60°C 30sec, 72°C 30sec], 72°C 5min) and run on a 2 % agarose gel (Fig. S9).

### Tissue disaggregation

Tissue was disaggregated using the Omni-ATAC-seq protocol ^1^ with minor modifications. All steps were performed on ice or in a 4°C chilled centrifuge. Briefly, mice were sacrificed and cortices were rapidly dissected and transferred into a chilled 7 mL Dounce Homogenizer containing 2.5 mL HB (320 mM sucrose, 0.1 mM EDTA, 0.1 % NP40, 5 mM CaCl2, 3 mM Mg(Ac)2, 10 mM Tris pH 7.4, protease inhibitors (Pierce #88666), 0.016 mM PMSF). Tissue was homogenized using a Dounce Tissue Grinder set (Wheaton #357542) with 10 strokes using a loose pestle and filtered through a 100 µm nylon mesh (VWR #10199-658) followed by 20 strokes using a tight pestle and centrifuged for 1 minutes at 100g. 2 mL of supernatant was mixed with 2 mL 50 % iodixanol solution (50 % iodixanol in 1x homogenization buffer). 2 mL of a 29 % iodixanol solution (29 % iodixanol in 1x HB containing 480 mM sucrose) was layered under the 25 % iodixanol/tissue mixture. 2 mL of a 35 % iodixanol solution (35 % iodixanol in 1X homogenization containing 480 mM sucrose) was layered underneath the 29 % iodixanol solution. Nuclei were centrifuged for 20 minutes at 3,000g in a swing-bucket rotor. Nuclei at the 29 % and 35 % interface were transferred to a new tube and diluted 1:5 in ATAC-RSB (10 mM Tris-HCl pH 7.4, 10 mM NaCl, and 3 mM MgCl2 in water) and centrifuged for 5 minutes at 500g. Nuclei were resuspended in 1X Tagmentation Buffer (10 mM Tris pH 7.4, 5 mM MgCl2, 10 % DMF, 33 % 1X PBS (without Ca++ and Mg++), 0.1 % Tween-20, 0.01 % Digitonin (ThermoFisher #BN2006) and counted using Trypan Blue on a hemocytometer.

### Tn5 Transposase

Tn5 was produced exactly as described ^2^ with no modifications. For Omni-ATAC-seq, Tn5 transposase was assembled using pre-Annealed ME-A and ME-B (Table S3). For sci-ATAC-seq, Tn5 transposomes were assembled using pre-Annealed ME-C and ME-D oligonucleotides (Table S3). Oligonucleotides were annealed in H2O by combining ME-A, ME-B, ME-C, or ME-D oligos at 25 µM to Tn5MErev and incubating for 2 minutes at 94°C followed by a 0.1°C/s ramp to 25°C. ME-A and ME-B hybridized oligos were combined in equal quantities at a final concentration of 2.5µM and incubated with Tn5 transposase at a final concentration of 1.625 µM. ME-C and ME-D hybridized oligos were each incubated separately at a final concentration of 2.5 µM with Tn5 transposase at a final concentration of 1.625 µM to make separated strip-tubes each containing a single barcode. The buffer in which hybridized Tn5 transposomes were stored consists of a mixture of 40 % glycerol and 43.5 % Tn5 dialysis buffer (100 mM HEPES-KOH pH 7.2, 200 mM NaCl, 20 mM EDTA, 2 mM DTT, 20 % Glycerol, 0.2 % Triton X-100 in DEPC H2O). Enzyme was stored at −80°C.

### sci-ATAC-seq

Sci-ATAC-seq was performed using a protocol based on snATAC-seq ^3^. Nuclei were resuspended at 600,000 cells/mL in 1X Tagmentation Buffer. 8 µL of nuclei were aliquoted to each well of a 96-well plate (4,800 cells/well). 1 µL of each ME-C or ME-D carrying barcoded transposome at ∼1.5 µM was added to each well and gently vortexed. Tagmentation was performed at 37°C for 1 hour, briefly vortexing once at 30 minutes. 10 µL of 40 µM EDTA was added each well, briefly vortexed, and incubated at 37°C for 15 minutes to inactivate the Tn5. 20 µL sort buffer (2 % BSA and 2 mM EDTA in PBS (without Ca++ and Mg++)) was added to each well. Nuclei from each well were pooled, filtered through a 35 µm mesh (Corning #352235) and Draq7 (Abcam #ab109202) was added to a final concentration of 3 µM (35-38 µL). 25 single nuclei were sorted into each well of a 96-well plate using a BD FACSAria Fusion and transferred to a −80°C freezer.

Frozen plates containing sorted nuclei were thawed on ice. 2 µL of 0.2 % SDS was added to each well and incubated for 7 minutes at 55°C. 2.5 µL of 10 % Triton X-100 was added to each well. 2 µL of 25 µM Primer i5 and 2 µL of 25µM Primer i7 was added to each well. PCR was performed for using Q5 DNA polymerase (NEB #M0491S) with 1X GC buffer (72°C 5min, 98°C 30sec, 15 cycles of: [98°C 10sec, 63°C 30sec, 72°C 30sec], 72°C 5min). In order to minimize batch effect, all PCRs were performed sequentially on a single machine. All wells were then pooled and diluted 5:1 (∼24 mL) in Buffer PB (Qiagen #19066) with 1/20^th^ volume of 3 M NaOAc pH 5.2 (∼1.2 mL). This solution was run through MinElute columns (Qiagen #28004) using a QIAvac apparatus (Qiagen #19413) and washed once using 750 µL Buffer PE (Qiagen #19065). Samples were eluted twice using EB (10 mM Tris pH 8) warmed to 55°C. Samples were size selected using a 0.5X Ampure XP cleanup to remove large fragments followed by 3 consecutive 1.5X Ampure XP (Beckman Coulter # A63880) bead cleanup followed by a final 1.2X Ampure XP cleanup to completely remove all residual primers and resuspended in a final volume of 20 µL EB using the manufacturer’s recommended protocol.

### Omni-ATAC-seq

Tagmentation was performed using the Omni-ATAC-seq protocol ^1^ with modifications to inactivation and size selection. Briefly, 100,000 nuclei were tagmented in parallel to sci-ATAC-seq in 1X Tagmentation Buffer (10 mM Tris pH 7.4, 5 mM MgCl2, 10 % DMF, 33 % PBS (without Ca++ and Mg++), 0.1 % Tween-20, 0.01 % Digitonin) using 100 nM of ME-A and ME-B bound Tn5 Transposase for 30 minutes at 37°C. Tagmentation was inactivated with the addition of 5 volumes SDS Lysis Buffer (100mM Tris pH 7.4, 50 mM NaCl, 10 mM EDTA, 0.5 % SDS in DEPC H2O) and 100 µg Proteinase K (Invitrogen #25530049) for 30 minutes at 55°C followed by Isopropanol Precipitation using GlycoBlue (Invitrogen #AM9516) as a carrier. Samples were resuspended in EB and size selected using a 0.5X Ampure XP cleanup to remove large fragments followed by a 1.8X Ampure XP cleanup using the manufacturer’s recommended protocol. PCR was performed for using Q5 DNA polymerase (NEB #M0491S) with 1X GC buffer (72°C 5min, 98°C 30sec, 11 cycles of: [98°C 10sec, 63°C 30sec, 72°C 30sec], 72°C 5min) followed by a final 1.8X Ampure XP cleanup.

### Cell-culture sci-ATAC-seq

Briefly, human A549 cells and mouse Hepa1-6 cells were quickly thawed at 37°C from tubes containing 1X freezing medium and resuspended in 10 mL room temperature PBS (without Ca++ and Mg++). Cells were spun at 500g for 5 minutes at room temperature, resuspended in PBS, and counted using Trypan Blue on a hemocytometer. 450,000 A549 cells were mixed with 450,000 Hepa1-6 cells and lysed via the Omni-ATAC-seq protocol in 50 µL ice-cold ATAC-RSB-Lysis buffer (10 mM Tris pH 7.4, 10 mM NaCl, 3 mM MgCl2, 0.1 % NP-40, 0.1 % Tween20, 0.01 % Digitonin in DEPC H2O). Nuclei were washed in 1 mL (10 mM Tris pH 7.4, 10 mM NaCl, 3 mM MgCl2, 0.1 % Tween20) and spun at 500g for 10 minutes. Nuclei were then processed identically to our sci-ATAC-seq protocol with no modifications. Reads were counted at each barcode in either the mm10 or hg19 genome. Any cell containing both human and mouse reads was identified as a collision. Collision rate was calculated as 2*(collisions/total cells) ^4^.

### Library quality control, quantification, and sequencing

Library fragment distribution was measured using a bioanalyzer for nucleosome patterning. Following this, libraries were subjected to digital PCR ^5^ on a Bio-Rad QX200 droplet digital PCR system using oligos from Table S3. Libraries were loaded at 8pM on a Nextseq500 mid lane PE 150bp. Sequencing was performed using a custom recipe for the following, read 1: [36 imaged cycles], Index 1: [8 imaged cycles, 27 dark cycles, 8 imaged cycles], Index 2: [8 imaged cycles, 21 dark cycles, 8 imaged cycles], Read 2: [36 imaged cycles].

### Preprocessing and alignment

Libraries were preprocessed and aligned as in Preissl et al ^3^ with minor modifications. Briefly, read names were labeled with combined 32bp index reads. Pair-end reads were aligned to mm10 (or mm10 and hg19 for collision tests) using Bowtie2 in pair-end mode with parameters, -p 5 -t ­X2000–no-mixed–no-discordant and reads < MAPQ 30 and flag=1804 were removed and deduplicated. Barcode arrangement and orientation is altered in our Nextseq500 run compared to the HiSeq run used in Preissl et al. Each 8bp Barcode was corrected within a 2bp edit distance to their nearest barcode and separated into individual cells based on their barcode combination after which PCR duplicates and mitochondrial were removed. Cells were filtered by a minimum of 20 % fraction of reads in peaks (FRiP) score and an 820 read per cell cutoff. Peaks were called using MACS2 ^6^ with parameters, -g mm -p 0.05 --nomodel --shift 150 --keep-dup all (Table S4). Reads were assigned to RefSeq ^7^ gene bodies flanked by 2kb on each side or to MACS2-called peaks using Rsubread featurecounts ^8^. Insert size metrics were plotted using ATACseqQC ^9^ using pseudobulks from aggregated samples. Copy number analysis was performed using HMMcopy as in Knouse et al ^10^ with no modification (Figure S3, Table S5).

### Clustering

Features were counted at gene body tiles extended by 2kb using Rsubread Featurecounts, covering ∼65 % of reads per sample. Dimensional reduction was performed on the first 50 principal components using Monocle 3.0 Alpha ^11^ on the first 50 principal components on gene body reads using UMAP. Clusters were called using Louvain clustering (Table S2b). 3D plots were generated using plot3D ^12^. Cells were counted by dividing the number of cells from each replicate of each cluster by the total number of cells sequenced per replicate (% total) (Table S2c). Differential cell counts were visualized and compared using ggpubr ^13^. Clustering of the top 20,000 TF-IDF selected peaks in Figure S6 was performed as in Cusanovich et al ^14^ with no modifications.

### Cell-type identification

Psuedobulk bam files from called clusters were merged using Samtools. Features were called at gene body tiles extended by 2kb using Featurecounts. Reads were filtered for a CPM > 1 in at least 4 samples. Cell-types were clustered based on their relative average accessibility at marker genes. Cell-type-specific markers (p.adj < 0.05 log2FC > 1) were called using Limma topTreat ^15^by comparing to genes enriched in each cluster or cell-type compared to all other clusters (Table S6) or cell-types (Table S7). Plots for cell-type identification were generated using log2CPM values normalized per gene across all clusters on a scale of 0 to 1; average scaled values were used for visualizations. Relative accessibility was visualized using ggplot2. Clustering was performed using hclust and visualized using ggdendro ^16^.

### Differential accessibility, Gene ontology and disease ontology

Pseudobulk bam files from called clusters were combined using samtools. Features were called at all peaks using Featurecounts. Differential accessibility (p.adj < 0.05 and |log2FC| > 0.585) was called using Limma topTable (Table S7). Peaks were assigned to their nearest RefSeq gene; RefSeq files were downloaded from UCSC table browser ^17^. Gene ontology and disease ontology analysis was performed on differentially accessible gene bodies and peaks (Table S9) using ClusterProfiler ^18^ and DOSE ^19^ respectively using clusterCompare with fdr correction. Dotplots were visualized using ClusterProfiler. Heatmaps were visualized using pheatmap.

### Motif Analysis

Motif analysis was performed on differentially accessible regions using HOMER ^20^ against a background set of all called peaks (Table S10). HOMER data was aggregated using marge ^21^ for downstream analysis.

### Comparison to Omni-ATAC-seq

Omni-ATAC-seq samples single-end reads were aligned to mm10 using Bowtie2 in pair-end mode with parameters, -p 5 and reads < MAPQ 30 and flag=1804 were removed and deduplicated. Read were assigned to peaks called in our sci-ATAC-seq libraries using Featurecounts. Libraries were downsampled to approximately 3.3M assigned reads using metaseqR downsample.counts ^22^. Scatterplots and Pearson correlations were visualized LSD ^23^.

### Genome browser tracks

Bigwig files were generated using Deeptools 3.0.2 ^24^ bamCoverage --binSize 1 --normalizeUsing RPKM --ignoreForNormalization chrM using cell-clusters merged using from all samples and displayed using GViz ^25^.

