## Supplemental figures 1-9 for "Single cell ATAC-seq identifies broad changes in neuronal abundance and chromatin accessibility in Down Syndrome"

a. Representative FANS results

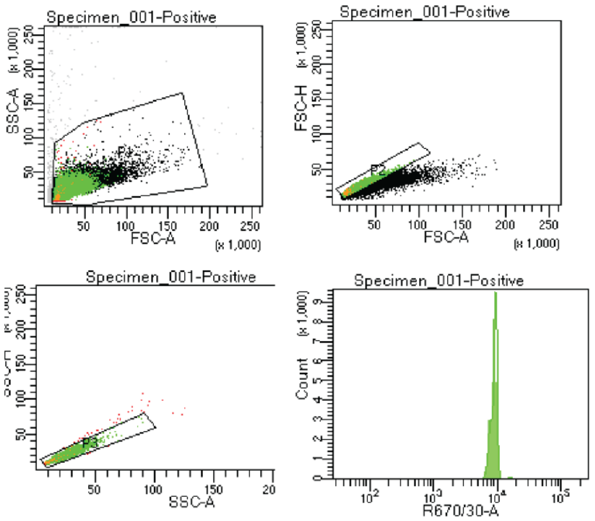


b. Bioanalyzer Trace

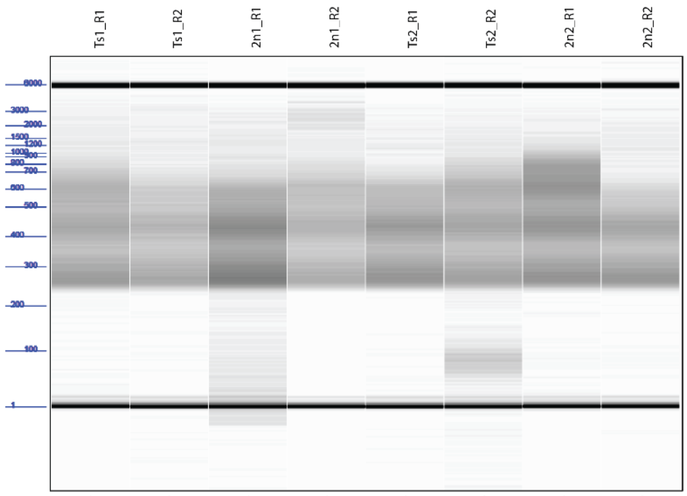


c. Representative ddPCR results

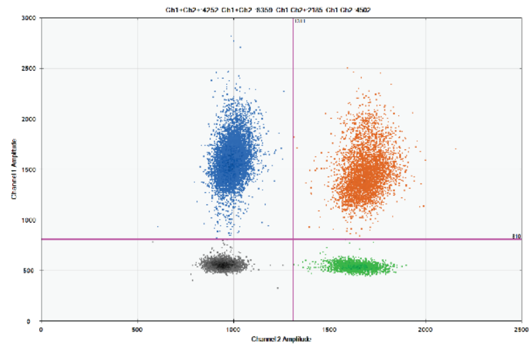

FAM (Ch1), custom library primers/probe: 1247 copies/uL
HEX (Ch2), reference primers/probe: 477 copies/uL
(1247 copies/uL) / ( 477 copies/uL) = 2.6
Known concentration of reference = 1 .1nM
2.6 * 1.1 = 2.9nM

d. FRiP cell in each assigned genotype

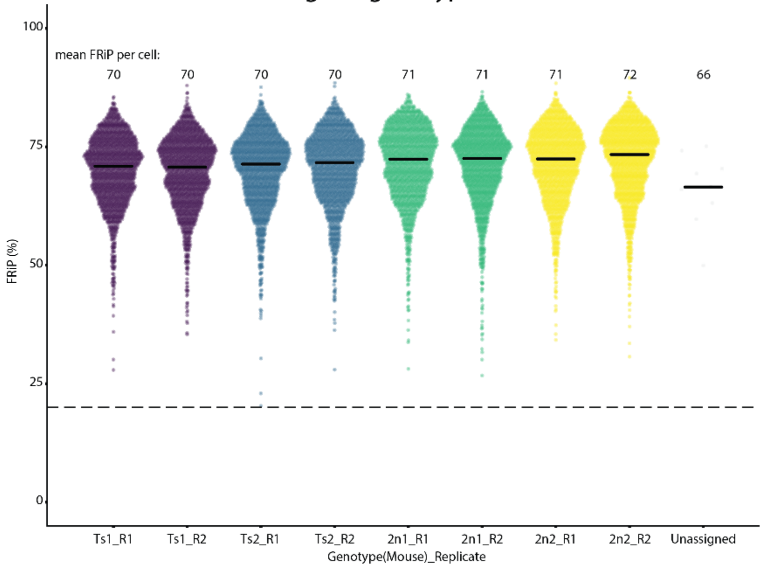


e. Fragment size distributions of Pseudobulks

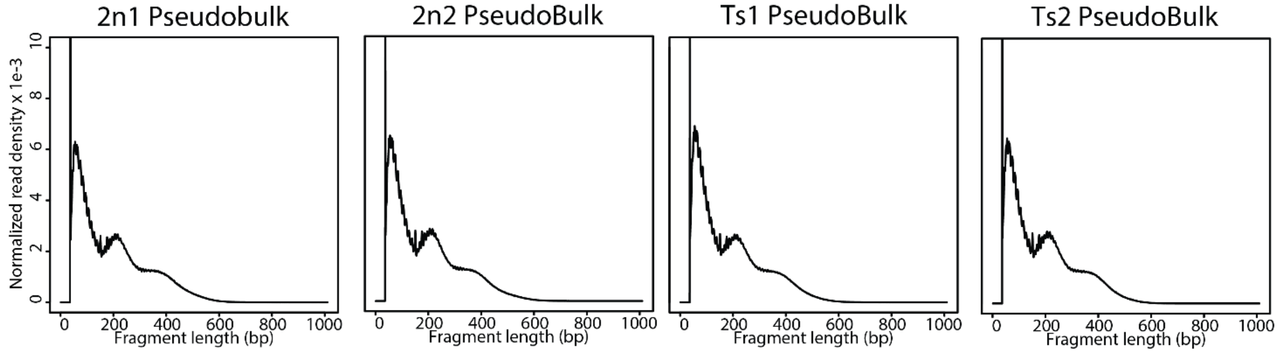

**Figure S1: Library QC. (a)** Representative FANS results showing FSC, SSC, and DRAQ7 gating (R670/30-A). **(b)** Bioanalyzer library sizes; figure has been globally contrast-adjusted for clarity. **(c)** Representative ddPCR results from a single library and control. **(d)** Distribution of FRiP per cells with reads > 820. **(e)** Fragment distribution of pseudobulks from QC-passed cells in each mouse.

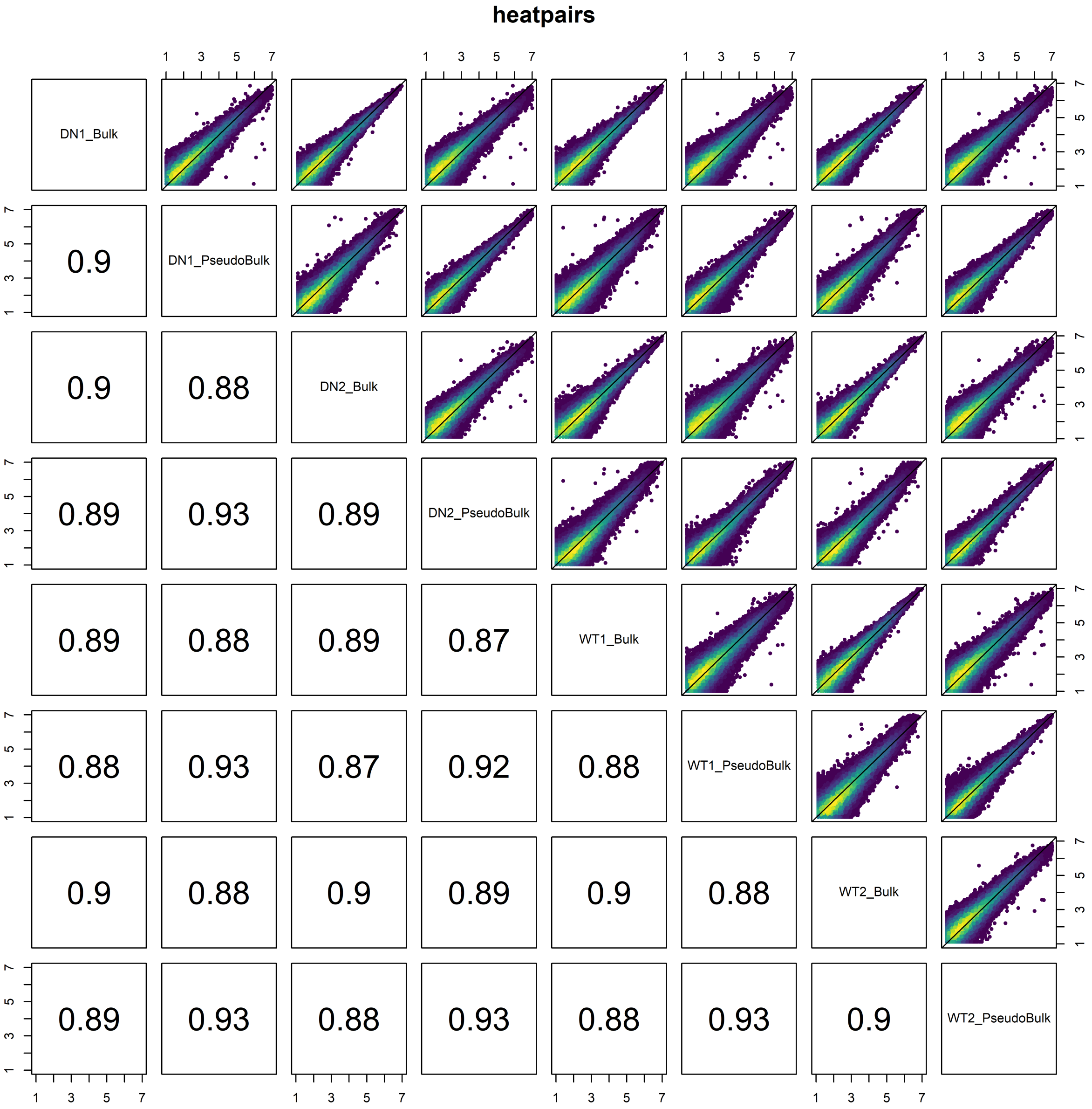


**Figure S2: Sample correlations.** Pearson correlation of all mice in bulk libraries (bulk) and pseudobulk libraries. Reads shown in log2CPM.

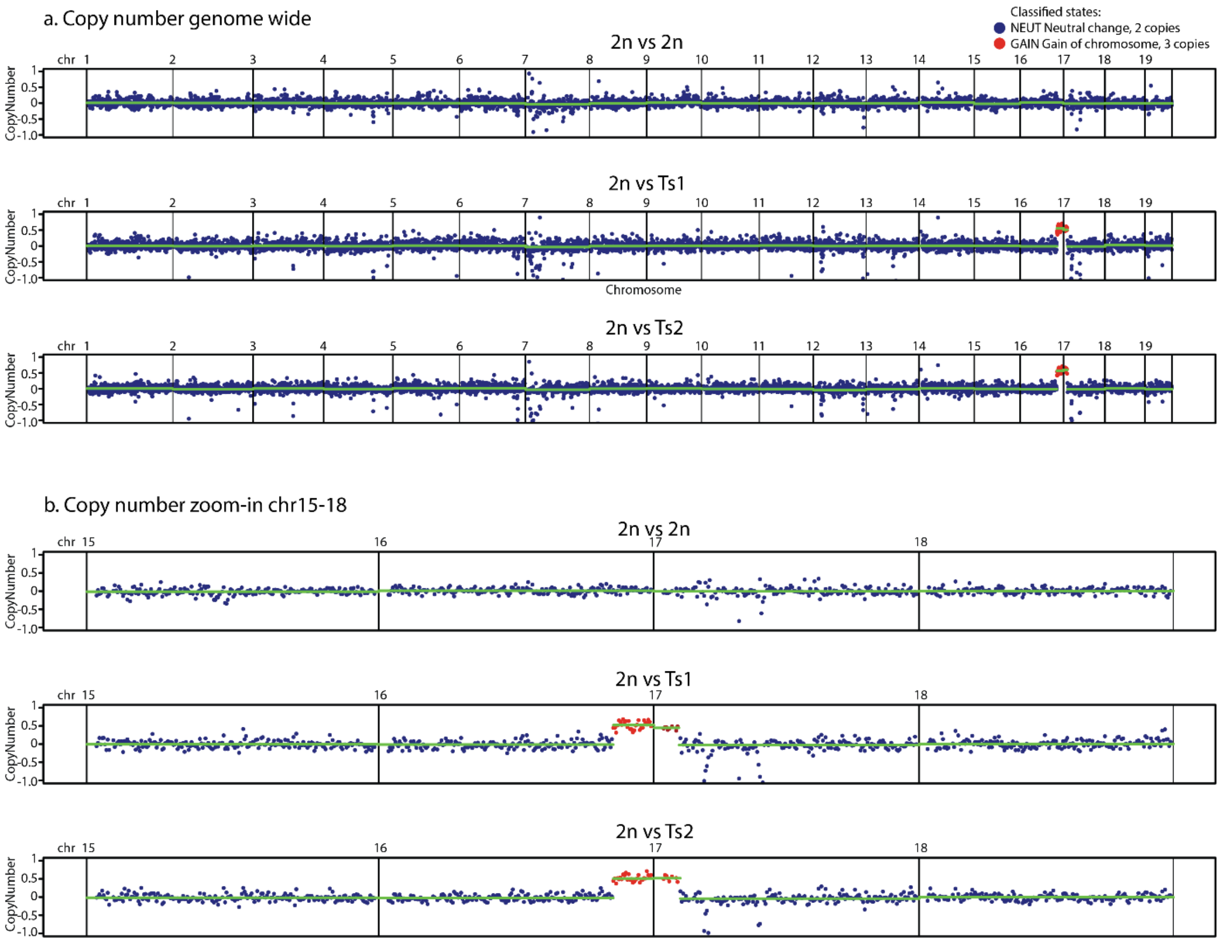


**Figure S3: Copy number confirmation**. (a) Copy number across genome compared to 2n animals at 500kb tiles using mm10. (b) Copy number zoomed in to chr15-18. Blue dots denote 2n state. Red dots denote 3n state at chr16:81,000,000-98,200,000 and chr17:3,500,001-9,000,000.

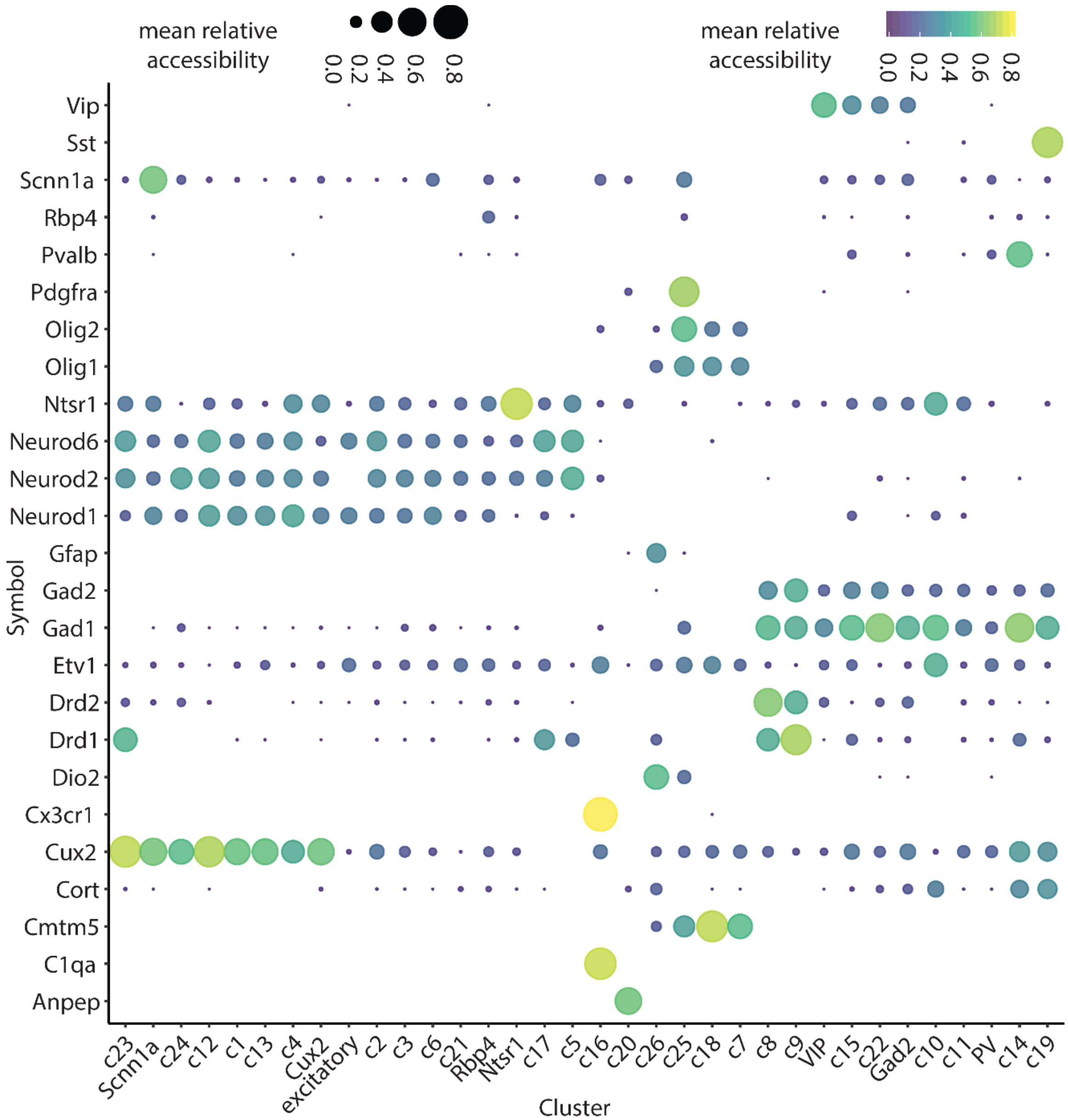


**Figure S4: Comparison to FACS-sorted data**. Cell-type specific markers at identified clusters and FACS sorted ATAC-seq data.

### S5a (top) S5b (bottom), cell-types:

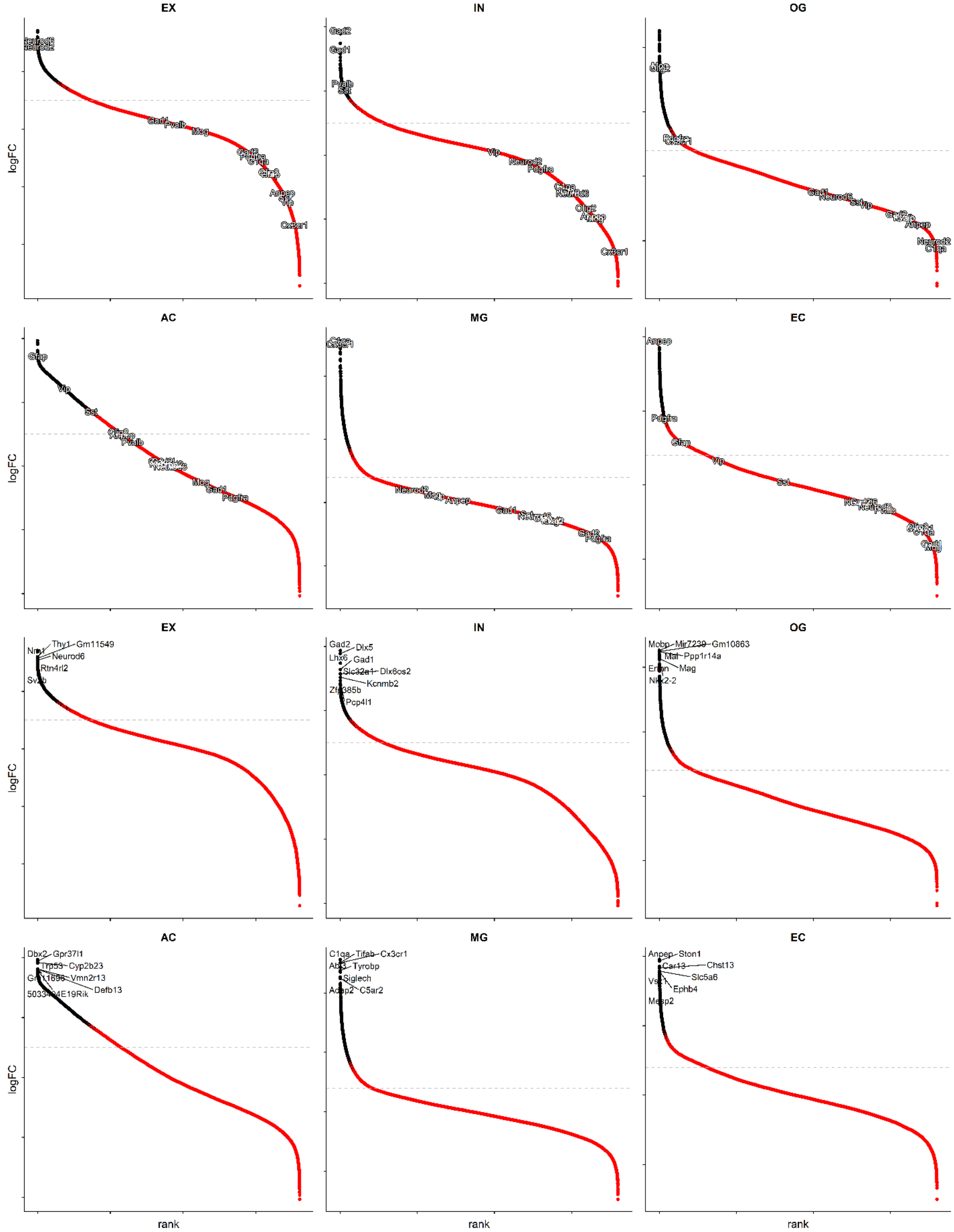

### S5c EX clusters:

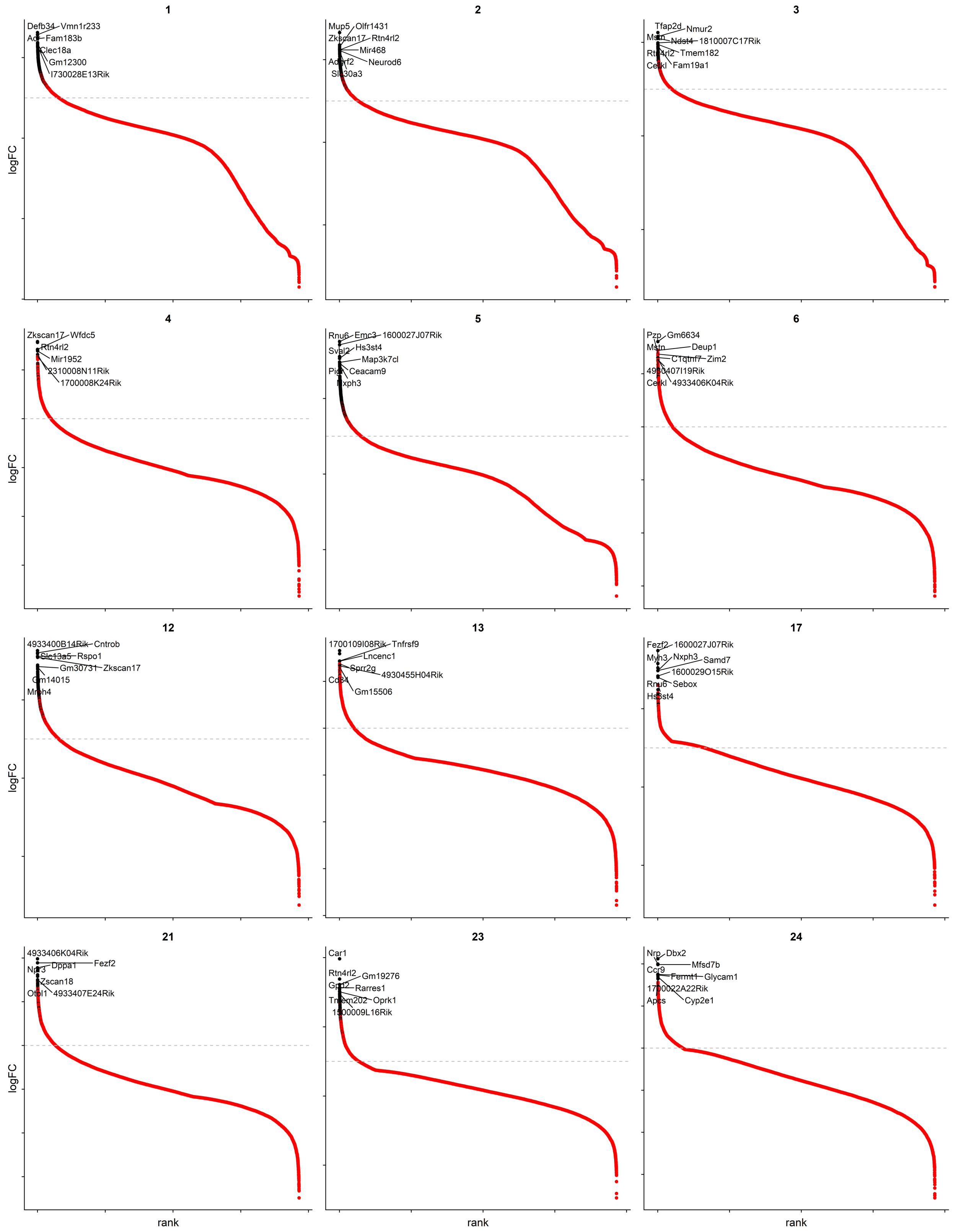

### S5d: IN clusters

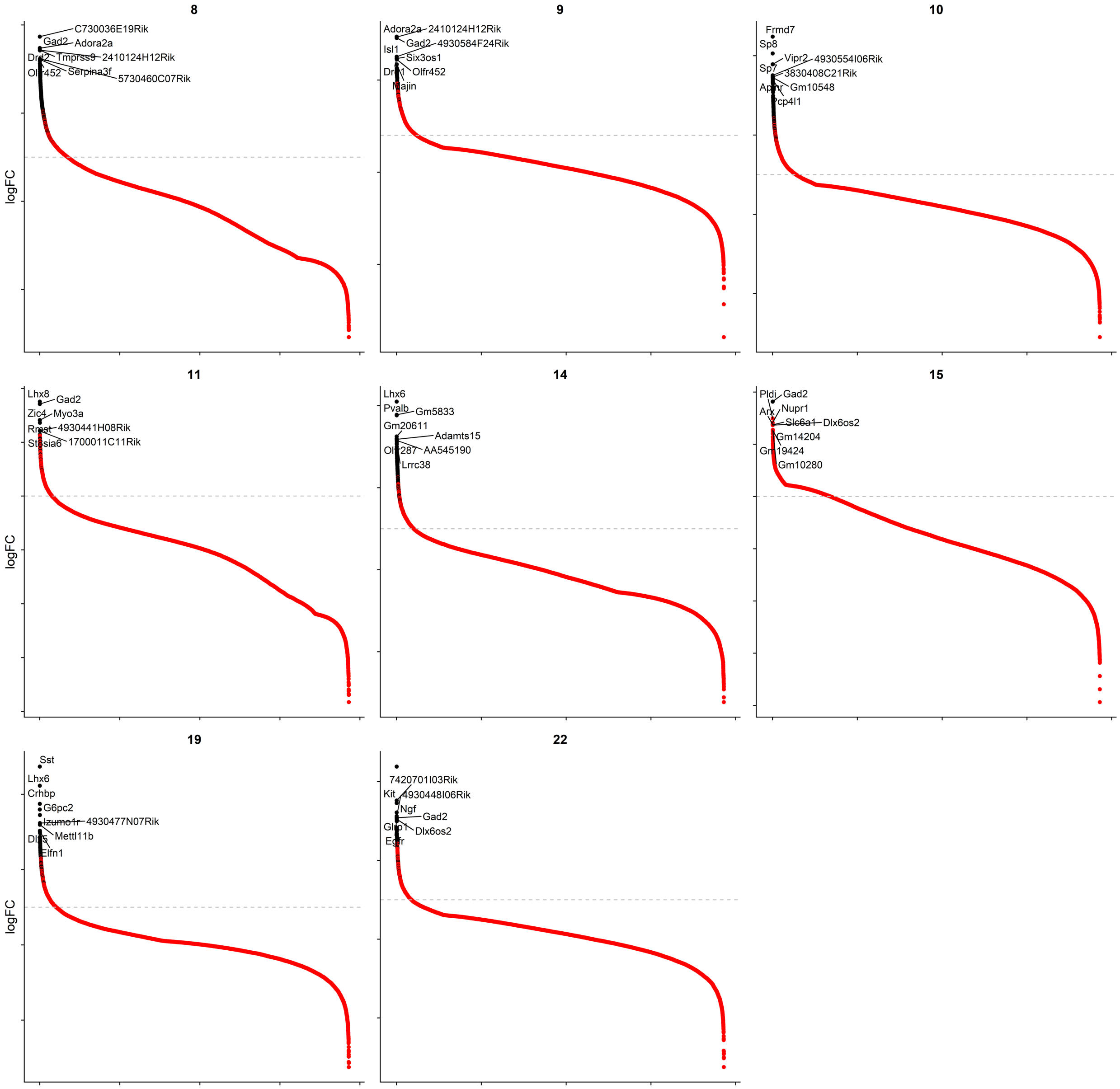

**S5e: glia clusters**

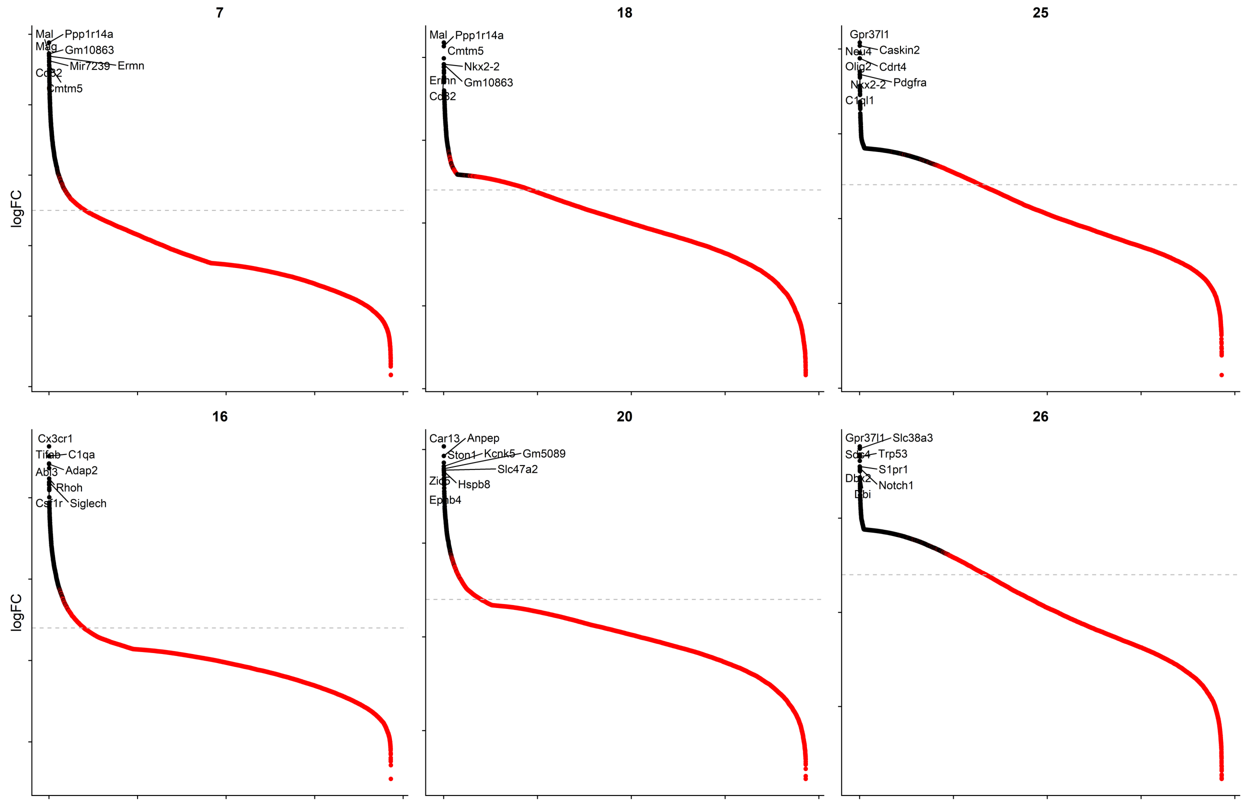


**Figure S5: Cell-type and cluster-enriched genes.** log2 Fold-change vs Rank at gene bodies in each cell-type showing **(a)** marker genes and **(b)** top 10 genes. Top10 hyper-accessible genes at each cluster sorted by EX (c), IN (d), and glia (e). Black dots denote p.adj < 0.05 and log2 fold-change > 1

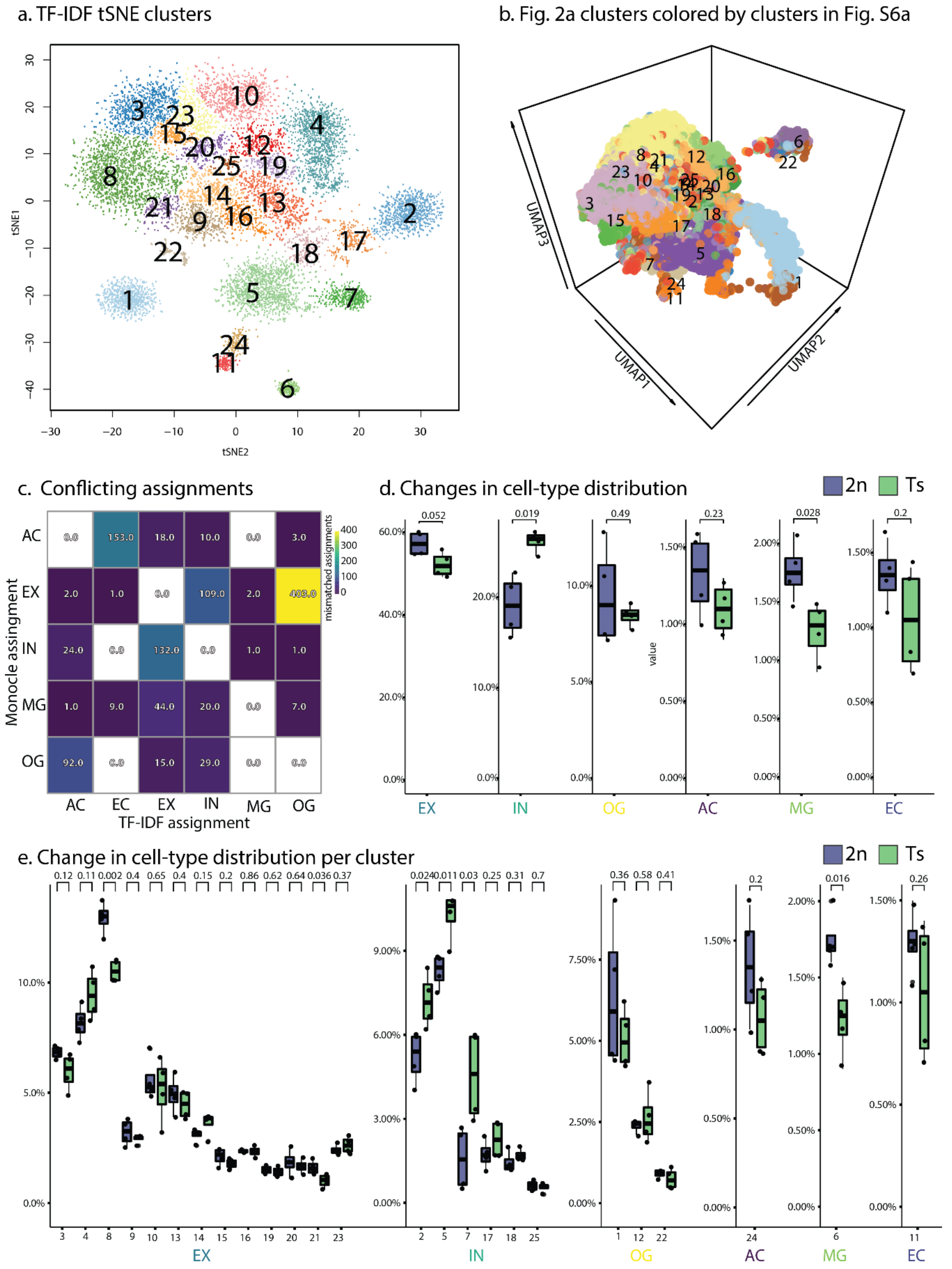


**Figure S6: Comparison to TF-IDF. (a)** t-SNE from 20,000 TF-IDF selected peaks. **(b)** Clusters in (a) overlaid on Fig. 2a UMAP. **(c)** Mismatched cell-type assignments. **(d)** Change in tell-type distribution. **(e)** Change in cell-type distribution per cluster. Blue bars denote 2n mice, green bars denote Ts mice.

a. Gene Ontology at differentially accesible peaks in Ts mice

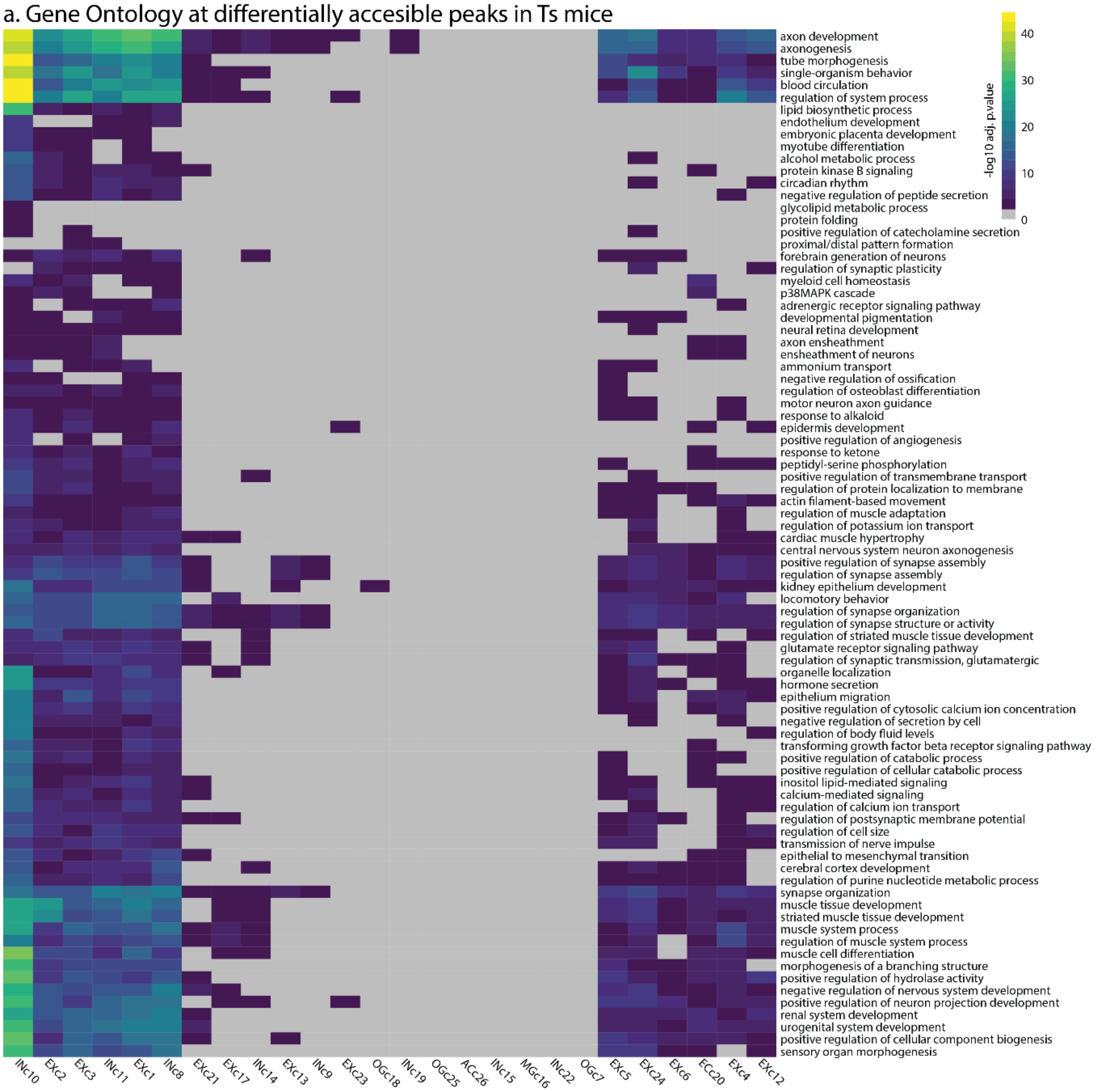


b. Disease Ontology at differentially accesible peaks in Ts mice

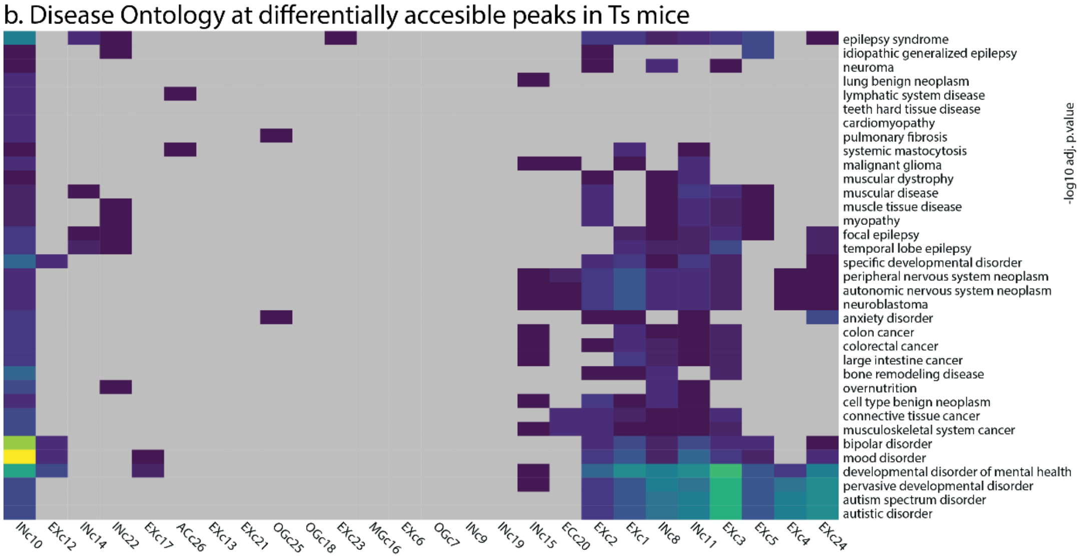

**Figure S7: Gene ontology and disease ontology changes per clusters. (a)** Gene ontology at nearest changed genes in Ts mice per cluster. Values denote fdr adjusted p. values filtered to a p value < 0.0001. **(b)** Disease ontology at nearest changed genes in Ts mice per cluster. Values denote fdr adjusted p. values filtered to a p value < 0.05.

### Motifs at differentially accessible peaks

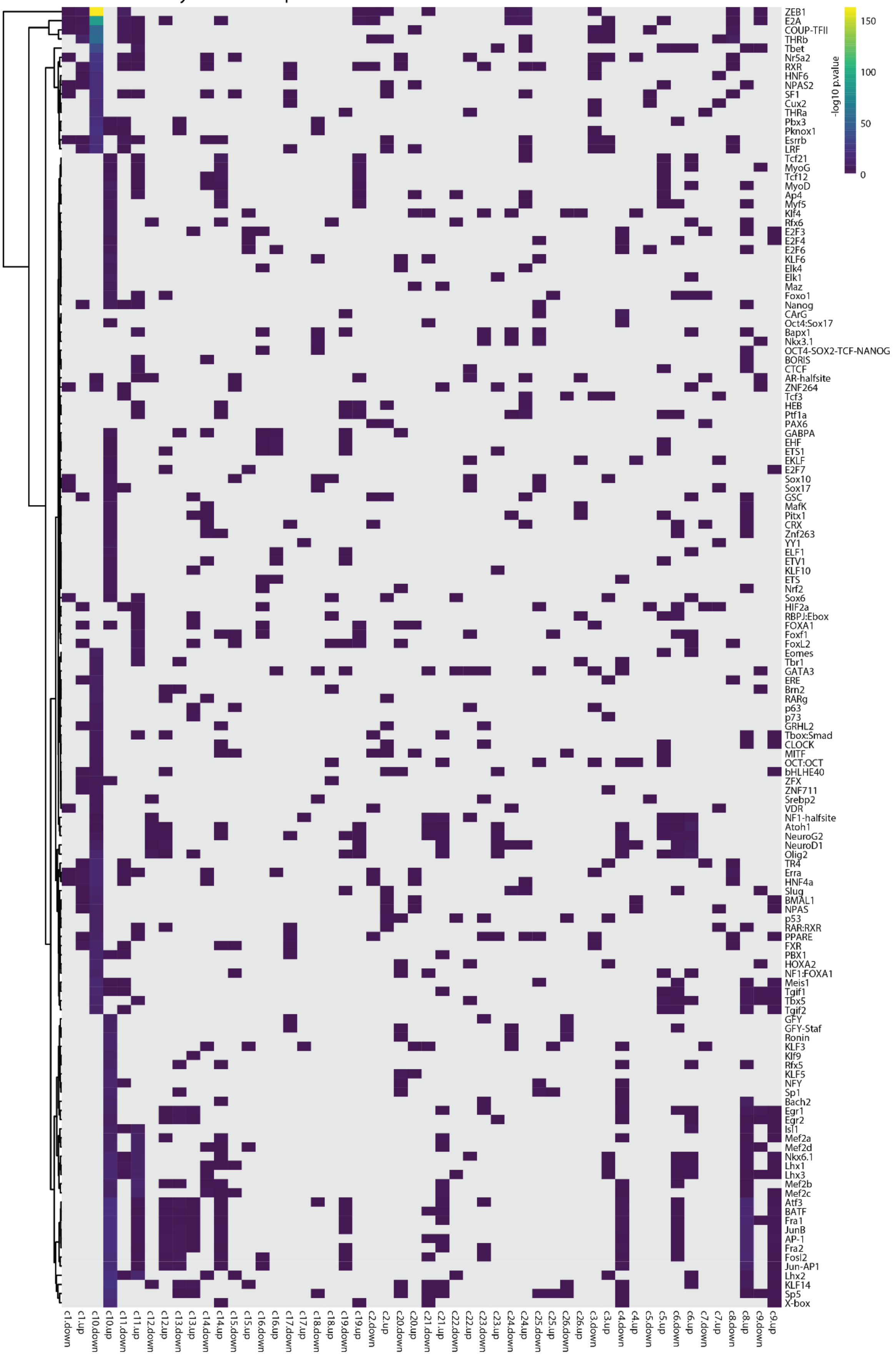

**Figure S8: Motifs at differentially accessible peaks in clusters.** Motifs changed at differentially accessible peaks in all clusters with a minimum p. value of 0.01 in at least one sample.

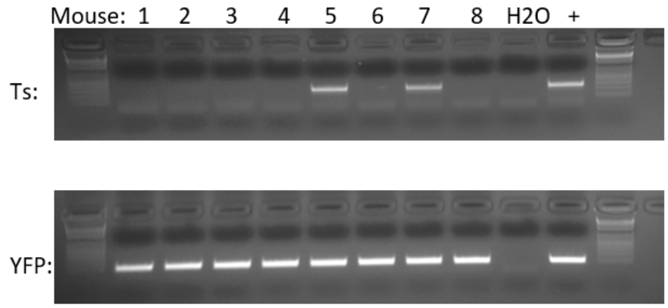


**Figure S9: Genotype.** PCR against the chr16-chr17 breakpoint in Ts65dn (top) and YFP (bottom) in Ts65Dn males used in sci-ATAC-seq. Mouse 5-8 were used in this study.
